# Theoretical analyses for the evolution of biogenic volatile organic compounds (BVOCs) emission strategy

**DOI:** 10.1101/2024.05.05.592540

**Authors:** Sotaro Hirose, Akiko Satake

## Abstract

Plants emit biogenic volatile organic compounds (BVOCs) as signaling molecules, playing a crucial role in inducing resistance against herbivores. Neighboring plants that eavesdrop on BVOC signals can also increase defenses against herbivores or alter growth patterns to respond to potential risks of herbivore damage. Despite the significance of BVOC emissions, the evolutionary rationales behind their release and the factors contributing to the diversity in such emissions remain poorly understood. To unravel the conditions for the evolution of BVOC emission, we developed a spatially-explicit model that formalizes the evolutionary dynamics of BVOC emission and non-emission strategies. Our model considered two effects of BVOC signaling that impact the fitness of plants: intra-individual communication, which mitigates herbivore damage through the plant’s own BVOC signaling incurring emission costs, and inter-individual communication, which alters the influence of herbivory based on BVOC signals from other individuals without incurring emission costs. Employing two mathematical models—the lattice model and the random distribution model—we investigated how intra-individual communication, inter-individual communication, and spatial structure influenced the evolution of BVOC emission strategies. Our analysis revealed that the increase in intra-individual communication promotes the evolution of the BVOC emission strategy. In contrast, the increase in inter-individual communication effect favors cheaters who benefit from the BVOCs released from neighboring plants without bearing the costs associated with BVOC emission. Our analysis also demonstrated that the narrower the spatial scale of BVOC signaling, the higher the likelihood of BVOC evolution. This research sheds light on the intricate dynamics governing the evolution of BVOC emissions and their implications for plant-plant communication.

## 1. Introduction

Plants emit large amounts of biogenic volatile organic compounds (BVOCs) into the atmosphere. BVOCs released from terrestrial ecosystems and plants are estimated to exceed 1,000 Teragram Carbon (TgC) per year globally and account for about 90% of total atmospheric VOC emissions (Guenther et al., 2012). BVOCs are lipophilic small molecules with high vapor pressure at normal temperature (Adebesin et al., 2017). Plants mainly release BVOCs such as terpenoids, fatty acid derivatives, benzenoids, and phenylpropanoids (Adebesin et al., 2017). The types of BVOCs and the composition of their blend exhibit considerable variability depending on the presence or absence of feeding damage, the growth stage of the plant, and environmental conditions (Hagiwara et al., 2021; Holopainen and Gershenzon, 2010; Šimpraga et al., 2019). The emission of BVOCs increases under stress conditions such as high temperatures, intense light, herbivore feeding damage, and pathogen invasion (Holopainen and Gershenzon, 2010). BVOCs released into the atmosphere from terrestrial ecosystems have been demonstrated to influence solar radiation and rainfall by contributing to the formation and growth of secondary organic aerosols (Hallquist et al., 2009; Peñuelas et al., 2009; Peñuelas and Staudt, 2010; Satake et al., 2024), and also play a role in tropospheric ozone production.

Despite the significance of BVOCs released from plants in shaping Earth’s surface, atmosphere, and climate, the evolutionary reasons behind plants emitting BVOCs and the factors contributing to the diversity in BVOC emissions are not well known. The evolution of the BVOC emission strategy is based on a trade-off concerning the costs and benefits of BVOC production. Since BVOCs are derived from carbon acquired through photosynthesis, their release as a signal can be costly. In a study of seven Mediterranean woody species, about 0.51-5.64% of the carbon acquired through photosynthesis in summer was released as terpenes (Llusia and Penuelas, 2000). In *Populus* x *canescens*, the amount of carbon C used in the synthesis of BVOCs was 1.6% in mature leaves and 7.0% in young leaves (Ghirardo et al., 2011). The emission of BVOCs play a role in facilitating plant-plant communication, as well as plant-animal communication such as seed dispersers, pollinators, and predators of herbivores. In addition to the increased defense of the elicited plant, BVOCs can diffuse around the elicited plant and are perceived by neighboring plants. Plants that eavesdrop on BVOC signals can improve their performance by increasing defenses against herbivores or altering growth patterns to respond to potential risks or opportunities (Karban, 2021), referred to plant-plant communications. Some of the most commonly emitted BVOCs serve as communication molecules that trigger cellular responses in the receiver plants. A recent study shows that isoprene and *β*-caryophyllene sensed by Arabidopsis plants can induce resistance to pathogens (Frank et al., 2021). BVOCs induced by herbivore feeding are commonly referred to as herbivore-induced plant volatiles (HIPVs), encompassing terpenoids and Green Leaf Volatiles (GLVs).

BVOCs have short lifetimes due to their high reactivity with the atmosphere, and most BVOCs in the atmosphere decompose in a few hours or less (Atkinson and Arey, 2003). Due to such short lifetimes, it is expected that BVOCs have a relatively short effective distance as signaling agents. Effective distances for plant-plant communication have been reported in different plant systems, ranging from 0.6 m for sagebrush, *Artemisia tridentata* (Karban et al., 2006) and 5-7 m for beech, *Fagus crenata* (Hagiwara et al., 2021). It suggests that BVOCs can act as signaling agents between individuals at relatively short distances. This short effective range of BVOC signaling could affect the evolution of the BVOC emission strategy. Therefore, to unravel the conditions for the evolution of BVOC emission, investigations conducted within a framework that explicitly considers the spatial distribution of plants are needed.

Previously, BVOC emission has been modeled as secondary signals by undamaged plants that promote the recruitment of natural predators of herbivores (Kobayashi et al., 2006; Kobayashi and Yamamura, 2007, 2003). Further theoretical assessment of the evolutionary conditions of BVOC emission strategies might lead to an in-depth understanding of BVOC emission from the viewpoint of the public goods game (Smith and Schuster, 2019). Public goods refer to resources or benefits that are available to all members of a group, regardless of whether they contribute to their production or maintenance. These goods are often characterized by non-excludability, meaning that individuals cannot be easily excluded from accessing the benefits. Public goods situations arise when individuals within a population can choose to contribute resources or efforts towards a collective benefit that is shared by all members of the group. However, there is often a temptation for individuals to “cheat,” meaning they benefit from the public good without contributing to its production or maintenance. In the context of BVOC emission, BVOCs released from plants can be interpreted as public goods which can be exploited by cheaters, who benefit from the BVOCs released from neighboring plants without bearing the costs associated with BVOC emission.

To elucidate the conditions driving the evolution of BVOC emission, we developed a spatially-explicit mathematical model that formalizes the evolutionary dynamics of BVOC emission strategies. We decomposed the communication effects into two categories: intra- and inter-communication effects. When a specific part or branch of an individual plant is damaged by herbivory, the plant induces defense mechanisms not only at the damaged part but also initiates or primes the resistance of intact parts or branches. This phenomenon occurs due to the perception of BVOC signaling originating from the damaged area (Frost et al., 2008; Heil and Adame-Álvarez, 2010). This effect is defined as intra-communication effect. Inter-communication effect is modeled as the induction of defense to herbivory by neighboring plants that eavesdrop and respond to BVOC signals emitted by the damaged plants. Through mathematical analyses and computer simulations of the model, we predicted the following three points: intra-individual signaling is necessary for the evolution of BVOC emission. As the spatial scale of BVOC signaling expands from the nearest neighbor to a wider range, the likelihood for the evolution of BVOC emission strategy decreases. BVOC emitters and non-emitters coexist across a broad spectrum of parameter settings.

## 2. Methods

### 2.1 An evolutionary game model of BVOC emission: a lattice model

We assumed that conspecific plant individuals adopt either a BVOC emission (E) or a non-emission (N) strategy. Although BVOC emission strategies are not clear-cut, we simplified the situation by assuming that the plants emitting smaller amounts of BVOC emission compared to BVOC emitters are BVOC non-emitters. Therefore, we define BVOC emitters and BVOC non-emitters in relative terms, which can be interpreted as BVOC high and low emitters, respectively. Both BVOC emitters and non-emitters trigger defense mechanisms of damaged organs against herbivory, thereby inducing resistance to herbivore damage BVOCs released from damaged plant organs play a role in initiating or priming resistance of undamaged parts of the same plant against herbivory (Frost et al., 2008; Heil and Adame-Álvarez, 2010). This phenomenon is referred to as the intra-communication effect, denoted as *α*_1_. We consider that the intra-communication effect in BVOC emitters is greater than that of BVOC non-emitters due to the substantial quantity of BVOCs emitted by the former. Most individual possess the capacity to eavesdrop and respond to BVOC signals emitted by their counterparts. The extent of their responsiveness to these BVOC signals is measured as the inter-communication effect, denoted as *α*_2_. The range of values for *α*_1_ and *α*_2_ is 0 < *α*_1_, *α*_2_ < 1. A portion of the carbon assimilated through photosynthesis is allocated for the biosynthesis of BVOCs (Ghirardo et al., 2011; Llusià and Peñuelas, 2000). This allocation entails a cost associated with BVOC emission, designated as *c*.

Based on the above assumptions, we developed an evolutionary game model that predicts the conditions in which the BVOC emission strategy can evolve by reference to the previous study about antipredator signaling (Kuga et al., 2021). The effective spatial range of signaling is relatively small, primarily limited to the nearest neighbor, due to the high reactivity of BVOCs. In addition, the spatial range of seed dispersal is limited and forms a spatially clumped distribution of plant individuals. To focus on the effect of spatial structure on the evolution of the BVOC emission strategy, we first consider the lattice model in which plant individuals distribute in the lattice structure (Fig. 1a and b).

**Figure 1:**
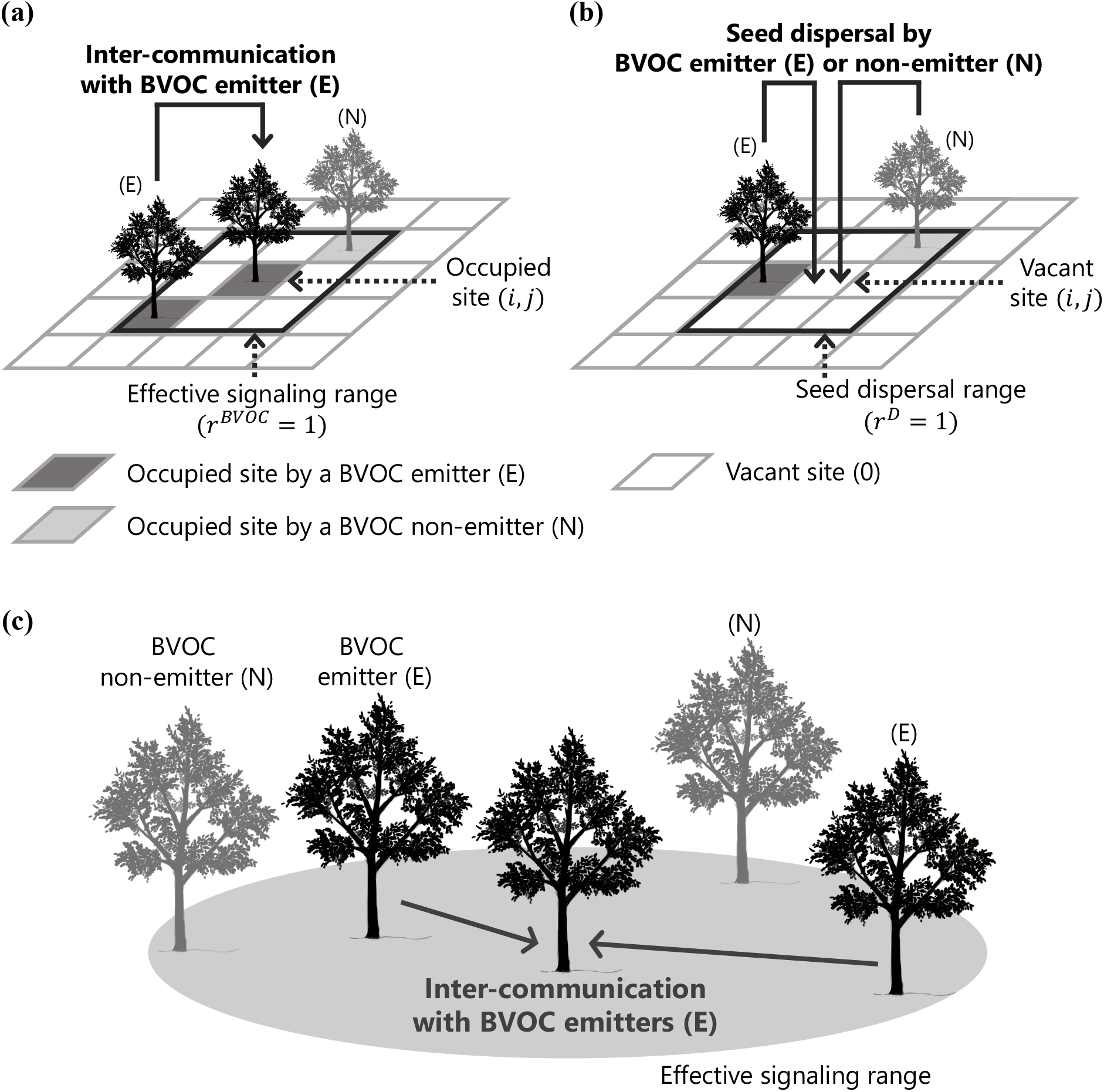
Examples of the inter-communication with BVOC emitters and the seed dispersal in the models. a, BVOC emitters (E) and non-emitters (N) can get the inter-communication effect via BVOC of nearby emitters within the effective signaling range (the black border area) in the lattice model. Lattice sites have three states: vacant (the white site), occupied by a BVOC emitter (the dark gray one), and occupied by a BVOC non-emitter (the gray one). The spatial range is defined by the distance *r*^*BVOC*^ from the central occupied lattice site (*i*,*j*); b, Individuals disperse their seeds to nearby vacant sites within the seed dispersal range (the black border area) in the lattice model. The range is defined by the distance *r*^*D*^ from the central vacant lattice site (*i*,*j*); and c, BVOC emitters and non-emitters can get the inter-communication effect via BVOC of *k* emitters (in Fig. 1c, *k* = 2) within the effective signaling range (the gray area) in the random distribution model.

In the lattice model, we evaluated how spatial structure affects the evolution of BVOC emission by introducing a two-dimensional square lattice model with a periodic boundary of size of 100 × 100. The state of the lattice site (*i*,*j*) is defined as follows:

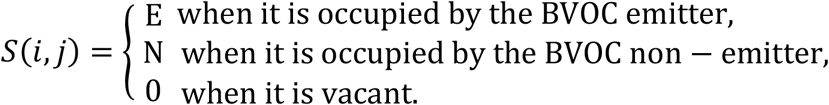

The state of each lattice site changes due to the mortality or recruitment of plant individuals. The recruitment rate is defined as the probability that each strategy occupies a vacant site, and this probability is affected by the fitness of each strategy.

The fitness of a plant individual is determined by the number of seeds produced by that individual, which depends on the herbivory damage and cost of BVOC emission. We assume that herbivory damage is mitigated by the intra- and inter-communication effects of BVOC. Let *p* denote the herbivory rate per unit of time when no intra- and inter-communication effects are at play. In the scenario of the BVOC emitter, the attenuation of herbivory damage is assumed to occur through the intra- and inter-communication effects of BVOCs. We formalize this mitigation effect as the product of the intra- and inter-communication effects, represented as 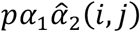. Here 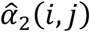 indicates the gross inter-communication effect of the individual at lattice site (*i*,*j*), which depends on the number of BVOC emitters within the neighborhood of *U*^*BVOC*^(*i, j, r*^*BVOC*^*E* with the range of *r*^*BVOC*^. Here we consider the Moore neighborhood of range *r* given as follows;

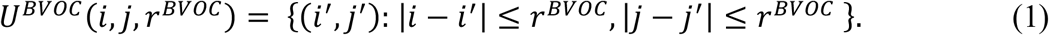

When *M* emitters exist within the neighborhood of *U*^*BVOC*^(*i, j, r*^*BVOC*^*E*, the gross inter-communication effect of the individual at the lattice site (*i*,*j*) is defined as follows:

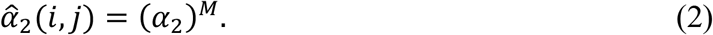

Conversely, BVOC emission incurs costs due to BVOC biosynthesis, thereby diminishing fitness by a factor of *c*. Given these assumptions, the fitness of the BVOC emitter at lattice site (*i*,*j*) is formalized as follows:

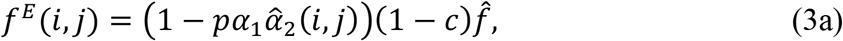

where 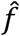 represents the number of seeds produced when there is no herbivory damage and BVOC emission. Given that the BVOC non-emitter does not gain any advantage from inter-communication without incurring costs, the fitness of the BVOC non-emitter at lattice site (*i*,*j*) is given as follows:

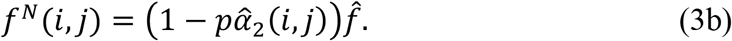

When the site is vacant, it generates no fitness (i.e. *f*^0^(*i, jE* = 0). The probability that the vacant site becomes occupied by the BVOC emitter or non-emitter depends on the relative fitness of the individuals included within the seed dispersal range. Let *F*(*i, j*) denote the total number of seeds dispersed to the (*i*,*j*) vacant site. *F*(*i, j*) is formalized as:

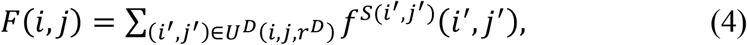

where *U*^*D*^(*i, j, r*^*D*^*E* is the neighborhood for seed dispersal of (*i*,*j*) site, given as follows;

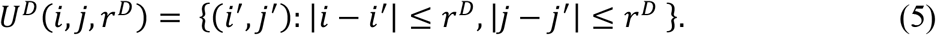

All parameters in the model are summarized in Table 1. The probability that the (*i*,*j*) vacant site becomes occupied by the BVOC emitters per each time step is:

**Table 1.** Variables or parameters and their definitions

| Variable/parameter | Definition |
| --- | --- |
| $x_E, x_N$ | Frequency of BVOC emitters and non-emitters |
| $n$ | Number of individuals within the signaling range |
| $k, M$ | Number of BVOC emitters within the signaling range |
| $\alpha_1$ | Effect of the intra-communication |
| $\alpha_2$ | Effect of the inter-communication with one individual |
| $\hat{f}$ | Number of seeds under the condition without herbivory |
| $p$ | Herbivore damage rate |
| $c$ | Cost of signaling |

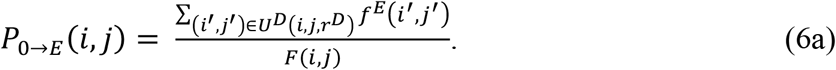

Similarly, the probability that the (*i*,*j*) vacant site becomes occupied by the BVOC non-emitters per each time step is:

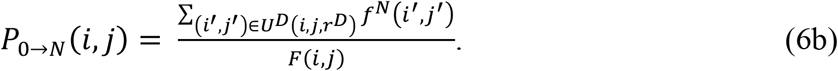

After the seed dispersal, BVOC emitters and non-emitters die and vacant sites are formed at the constant mortality rate of *d*. Thus, the transition rate from *E* to 0 (*P*_*E*→0_(*i, jE*) or *N* to 0 (*P*_*N*→0_(*i, jE*) at (*i*,*j*) site per each time step becomes *d*. We ran simulations with a parameter *d* = 0.0*D*. Three transition rates in the model are summarized in Table 2.

**Table 2.** Transition rates of a lattice site and its definitions

| Transition rate | Definition |
| --- | --- |
| $P_{0 \rightarrow E}(i, j)$ | Transition rate that the vacant site $(i, j)$ become occupied by a BVOC emitter per unit of time |
| $P_{0 \rightarrow N}(i, j)$ | Transition rate that the vacant site $(i, j)$ become occupied by a BVOC non-emitter per unit of time |
| $P_{E \rightarrow 0}(i, j), P_{N \rightarrow 0}(i, j)$ | Individual mortality rate per unit of time |

The spatial range of BVOC signaling and seed dispersal can largely affect the evolution of the BVOC emission strategy. We first consider the case in which *r*^*BVOC*^ = *r*^*D*^ = 1 in Eq. (1) both for BVOC signaling and seed dispersal equal. We then enlarged the spatial scales by considering all four possible combinations of the ranges for BVOC signaling and seed dispersal: (*r*^*BVOC*^, *r*^*D*^*E* = (1, 2), (2, 1), and (2,2). Our simulation is based on synchronized updating of the lattice state by repeating the following steps: (1) calculation of the fitness at each lattice site, (2) creation of vacant sites due to plant death, and (2) occupation of vacant site with the probability given in Eqs. (6a) and (6b). As an initial condition, we distributed a small number of BVOC emitters and non-emitters randomly in the lattice space filled with vacant sites. The combinations of the existence probabilities of them in the lattice space were as follows: (0.001, 0.001), (0.001, 0.0002), and (0.0002, 0.001). The fitness of individuals was determined by signaling effects with neighboring emitters based on the above assumptions. For a diverse parameter set, we ran independent simulations ten times over thirty thousand steps. To check whether emitters disappear, dominate, or coexist with non-emitters in equilibrium, we calculated the mean frequency of emitters in the lattice space filled with emitters and non-emitters at the thirty thousand steps before the occurrence of the stochastic mortality. For conducting simulations and numerical analyses in our model, we employed the Python programming language version 3.11.1, leveraging following key libraries: NumPy (v1.24.2) and Matplotlib (v3.7.1).

### 2.2 Approximating the spatially-explicit evolutionary dynamics of BVOC emission strategy by the random distribution model

Assuming that individuals are randomly distributed, we can formalize the evolutionary dynamics of the BVOC emission strategy by adopting the mean field approximation. We compared the results from the random distribution model to those from the lattice model to assess the effect of spatial distribution on the evolution of BVOC signaling. The spatial scale of BVOC signaling is defined as a circle expanding from the focal individual (Fig. 1c), and the expected number of the other conspecifics within the effective range of signaling is *n*. A higher value of *n* indicates that a population density is higher or the effective signaling range is larger. Let *P*(*k*) be the probability that *k* of *n* individuals emit BVOC in the effective signaling range. When plant individuals distribute randomly, *P*(*kE* follows a binomial distribution:

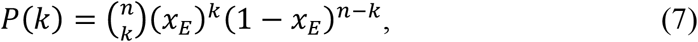

where *x*_*E*_ indicates frequency of BVOC emitters. When *n* BVOC emitters exist within the effective signaling range, the gross inter-communication effect is defined as follows:

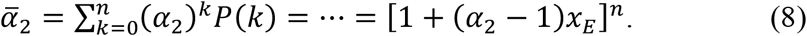

Subsequently, the fitness of BVOC emitters (*f*^*E*^) and non-emitters (*f*^*N*^) are given as:

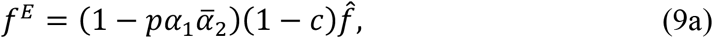

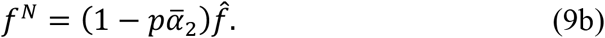

All parameters in the model are summarized in Table 1. Using the above equations, the evolutionary dynamics for the frequency of BVOC emitters *x*_*E*_ is formalized as:

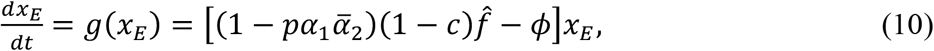

where *ϕ* indicates the population mean of fitness given as:

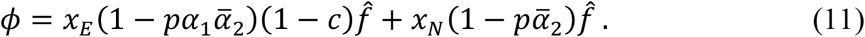

The range of parameters are expressed by inequalities: 0 < *p, c, α*_1_, *α*_2_ < 1. Since the equilibria *x*_*E*_^*^ satisfies *dx*_*E*_/*dt* = 0 (Table 3), we have:

**Table 3.** Equilibrium and the condition for an evolutionarily stable state (ESS)

| Value of $x_E^*$ | Type of equilibria | Condition for ESS |
| --- | --- | --- |
| $x_E^* = 1$ | Monomorphic | $0 < \alpha_1 < \frac{(\alpha_2)^n p - c}{(\alpha_2)^n p (1 - c)}$ |
| $0 < x_E^* < 1$ | Coexistent | $\frac{(\alpha_2)^n p - c}{(\alpha_2)^n p (1 - c)} < \alpha_1 < \frac{p - c}{p(1 - c)}$ |
| $x_E^* = 0$ | Monomorphic | $\frac{p - c}{p(1 - c)} < \alpha_1 < 1$ |

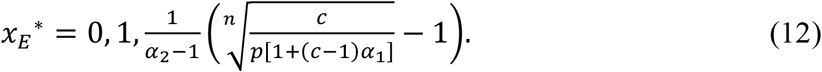

The equilibrium in which emitters and non-emitters coexist satisfies:

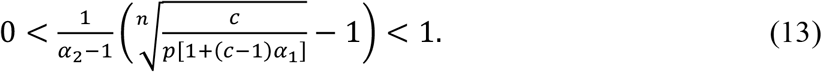

Since 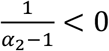 is satisfied, inequality (13) becomes:

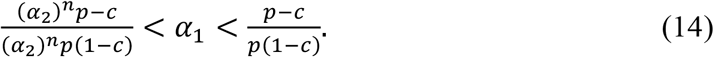

The parameter range where inequality (14) is satisfied on the *α*_1_ - *α*_2_ plane has a coexistent equilibrium. The other two equilibria are respectively monomorphic of non-emitters (*x*_*E*_^*^ = 0) or emitters (*x*_*E*_^*^ = 1). We evaluated the stability of these three equilibria by assessing the sign of *dg*/*dx*_*E*_, where *g* is formalized in Eq. (10). The equilibrium is locally stable if the sign of *dg*/*dx*_*E*_ is negative, which is the evolutionarily stable strategy (ESS).

The BVOC emission strategy evolves only when BVOC emission is advantageous even without an inter-communication effect. This condition is met when *f*^*E*^ > *f*^*N*^ with *α*_2_ = 1. This inequality becomes:

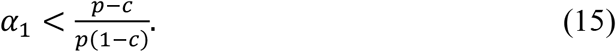

In the following, we analyze the case where 0 < *α*_1_ < (*p* − *cE*/(1 − *c*)*p* is satisfied. Moreover, as the random distribution model suggests that the BVOC emission strategy is more likely to evolve under the condition of the low BVOC production cost and the high herbivory rate (Appendix A), we will focus on such parameters (*c* = 0.1, *p* = 0.4) in subsequent analyses.

## 3 Result

### 3.1 The BVOC emission strategy is maintained when the intra-communication effect is strong but the inter-communication effect is weak

In the lattice model, we identified three parameter regions in the *α*_1_-*α*_2_ plane (Fig. 2a): ones where only BVOC emitters or non-emitters existed and another where both BVOC emitters and non-emitters coexisted. Plant individuals employing the BVOC emission strategy dominated the population when *α*_1_ is small and *α*_2_ is large, namely, when the intra-communication is effective compared to the inter-communication. The parameter region where only non-emitters existed (Fig. 2a) resulted from the stochastic extinction of emitters in the finite lattice structure with periodic boundary conditions.

**Figure 2:**
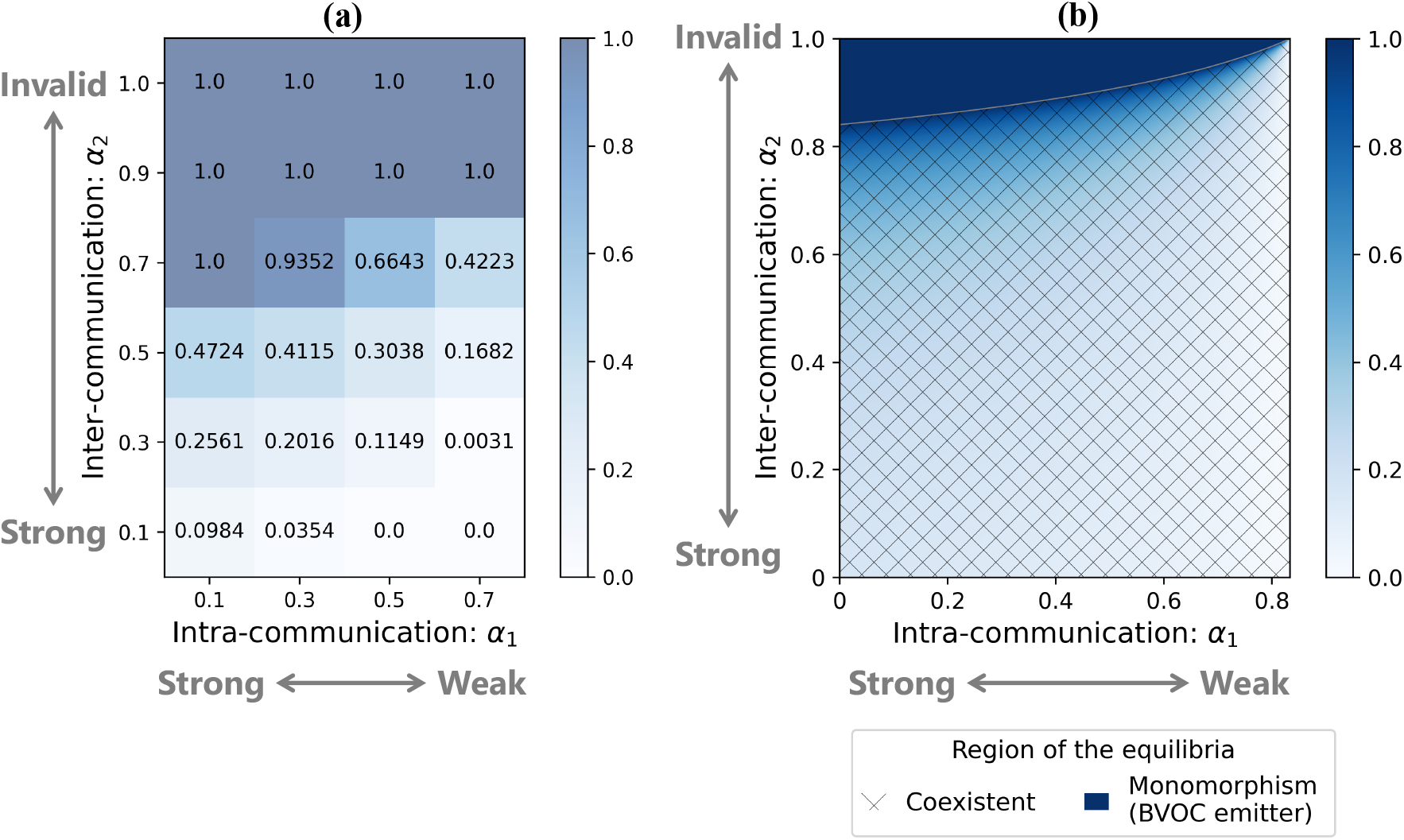
Effects of the intra- and inter-communication on the evolutionary consequences of the BVOC emission strategy. a, The average frequency of BVOC emitter obtained from simulations at thirty thousand steps in the lattice model. The color bar and decimals show the average frequency of emitters in ten times simulations. Parameters are *c* = 0.1, *p* = 0.4, *d* = 0.0*D, r*^*BVOC*^ = 1, *r*^*D*^ = 1 ; and b, Values of the local stable equilibria (the density of emitters) on the *α*_1_-*α*_2_ plane in the random distribution model. The parameter area can be divided into two types: The shaded area has a stable equilibrium where BVOC emitters coexist with non-emitters and the dark blue one has a stable equilibrium where BVOC emitters are dominant. Parameters are *c* = 0.1, *p* = 0.4, *n* = 8.

This finding was confirmed in the random distribution model. In the random distribution model, the BVOC emission strategy became dominant when intra-communication was more beneficial than inter-communication for mitigating herbivore damage (Fig. 2b). In most parameter regions, the BVOC emission strategy and the non-emission strategy coexisted, with the density of the BVOC non-emission strategy increasing as intra-communication effects rose. These results suggest that the BVOC emission strategy is more likely to evolve when intra-communication is more beneficial than inter-communication for mitigating herbivore damage.

### 3.2 The spatial structure affects the evolution of the BVOC emission strategy

We assessed the impact of the spatial range of BVOC signaling and the seed dispersal range on the evolution of the BVOC emission strategy using the lattice model. When the signaling range was limited to the neighboring trees (i.e., *r*^*BVOC*^ = 1), the density of BVOC emitters was higher compared to the case when it was wider (i.e., *r*^*BVOC*^ = 2) (Fig. 3a, 3b). This result suggests that the BVOC emission strategy is more likely to evolve when the spatial range of BVOC signaling is limited due to effective suppression of cheaters.

**Figure 3:**
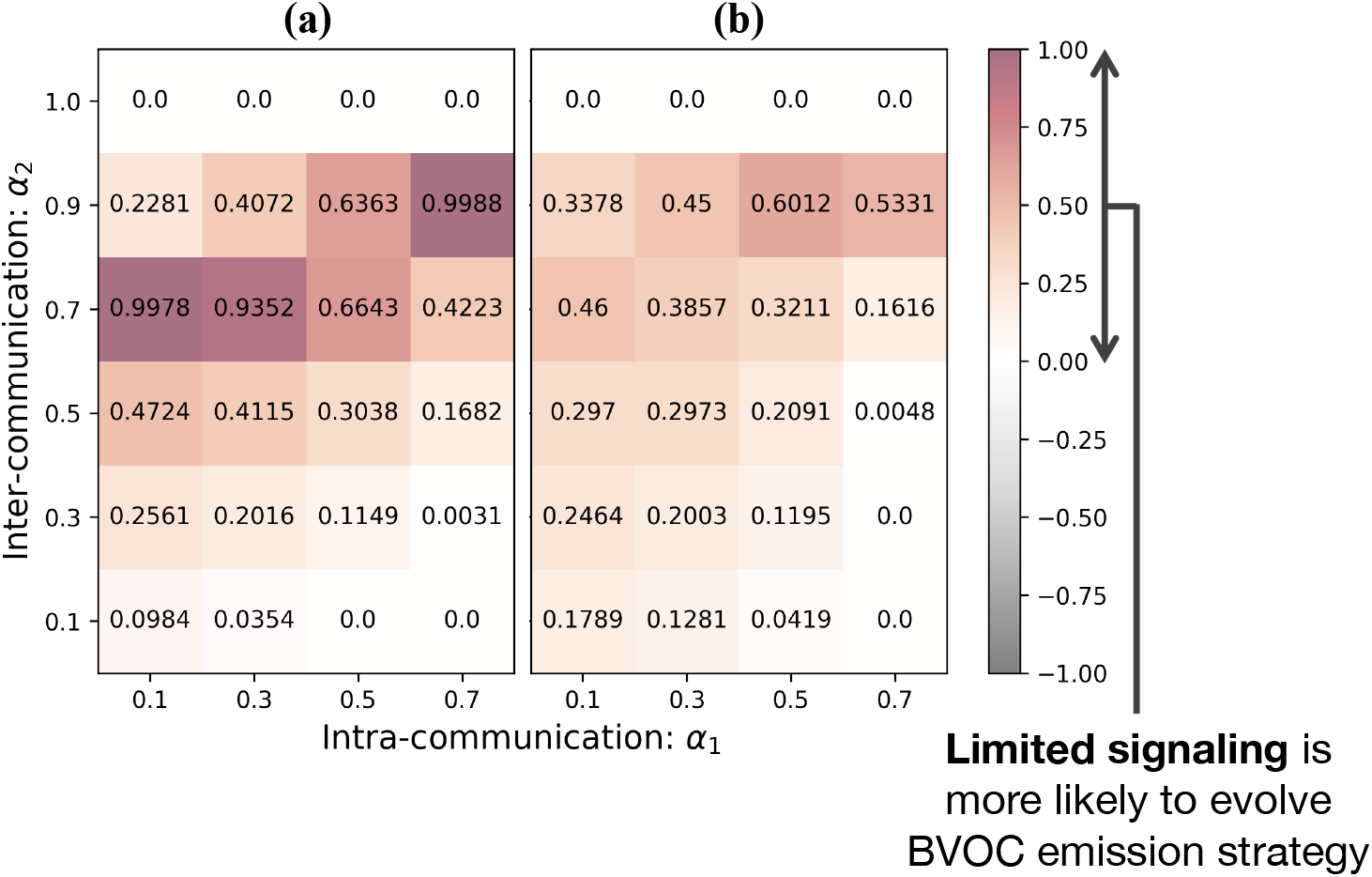
Effects of the spatial ranges of BVOC signaling on the evolutionary consequences of the BVOC emission strategy. a, The calculated difference in the average frequency of BVOC emitters in case the seed dispersal range is limited (*r*^*D*^ = 1) caused by different signaling ranges (*r*^*BVOC*^ = 1 and *r*^*BVOC*^ = 2). The red area, in contrast with the grey one, shows that the short signaling range (*r*^*BVOC*^ = 1) is advantageous for the fitness of the emission strategy. Parameters are *c* = 0.1, *p* = 0.4, *d* = 0.0*D* ; b, The calculated difference in the average frequency of BVOC emitters in case the seed dispersal range is wide (*r*^*D*^ = 2) caused by different signaling ranges (*r*^*BVOC*^ = 1 and *r*^*BVOC*^ = 2). Parameters are *c* = 0.1, *p* = 0.4, *d* = 0.0*D*.

The frequency of BVOC emitters was also affected by the spatial range of seed dispersal (Fig. 4a). When the seed dispersal range was limited, BVOC emitter formed clumped distributions (Fig. 4b, 4c). When the signaling range was limited (i.e., *r*^*BVOC*^ = 1) and the intra-communication is effective compared to the inter-communication, limited seed dispersal (i.e., *r*^*D*^ = 1) made the density of BVOC emitters higher compared to the case when the dispersal range was wider (i.e., *r*^*D*^ = 2) (Fig. 4a). These findings indicate that the clumped distribution of BVOC emitters plays a pivotal role in driving the evolution of BVOC emission strategies.

**Figure 4:**
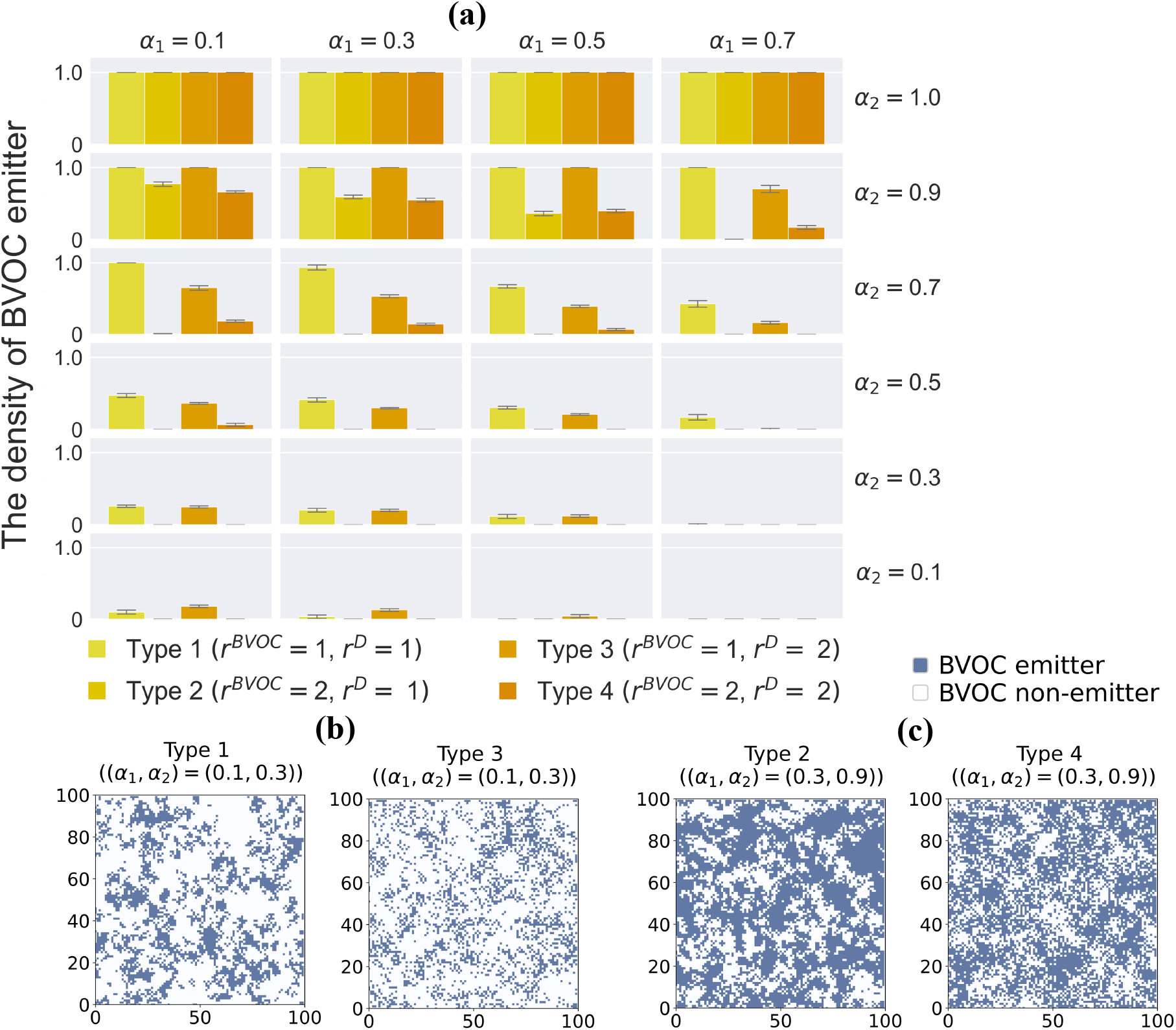
Effects of the spatial ranges of BVOC signaling and seed dispersal on the evolutionary consequences of the BVOC emission strategy. a, The average frequency of BVOC emitters of four combinations based on different signaling ranges (*r*^*BVOC*^ = 1 and *r*^*BVOC*^ = 2) and different seed dispersal ranges (*r*^*D*^ = 1 and *r*^*D*^ = 2). The yellow or orange bars correspond to the combinations of two ranges respectively, and the error bars show the standard deviation. Parameters are *c* = 0.1, *p* = 0.4, *d* = 0.0*D*; b, The examples comparing the distribution of individuals on the lattice structure at thirty thousand steps between simulations based on different seed dispersal ranges (*r*^*D*^ = 1 and *r*^*D*^ = 2). We ran simulations in case the signaling range is limited (*r*^*BVOC*^ = 1). The two figures respectively reflect the lattice space filled with BVOC emitters (blue) and non-emitters (white) at thirty thousand steps before the occurrence of the stochastic mortality of individuals, so there is no vacant site. Parameters are *c* = 0.1, *p* = 0.4, *d* = 0.0*D, α*_1_ = 0.1, *α*_2_ = 0.3 ; c, The examples comparing the distribution of individuals on the lattice structure at thirty thousand steps between simulations based on different seed dispersal ranges (*r*^*D*^ = 1 and *r*^*D*^ = 2). We ran simulations in case the signaling range is wide (*r*^*BVOC*^ = 2). Parameters are *c* = 0.1, *p* = 0.4, *d* = 0.0*D, α*_1_ = 0.3, *α*_2_ = 0.9.

## 4 Discussion

Despite the importance of BVOCs released from plants, the evolutionary rationales behind plants emitting BVOCs and the factors contributing to the diversity in BVOC emissions have remained elusive. To unravel the conditions for the evolution of BVOC emission, we developed a spatially-explicit mathematical model that formalizes the evolutionary dynamics of BVOC emission and non-emission strategies. Here, we have three important findings.

Our first main finding highlights the impact of intra-individual communication as a prerequisite for the evolution of the BVOC emission strategy. Secondly, we found that the narrower the spatial scale of BVOC signaling, the higher the likelihood of the evolution of BVOC emission. Lastly, we show the coexistence of both BVOC emitters and non-emitters across a broad spectrum of parameter settings. Regarding the first point, our theoretical finding aligns with the hypothesis that the release of BVOCs may have evolved to coordinate defenses within a plant rather than between plants (Heil and Adame-Álvarez, 2010). BVOCs released from damaged plant organs play a crucial role in priming unaffected parts of the same plant for the expression of resistance (Frost et al., 2008; Heil and Adame-Álvarez, 2010). Neighboring plants may be eavesdropping on cues that help them to prepare for upcoming danger (Frost et al., 2008). In this context, BVOCs released from plants can be interpreted as public goods (Smith and Schuster, 2019). Public goods are susceptible to exploitation by cheaters, who benefit from the BVOCs released from neighboring plants without bearing the costs associated with BVOC emission. Our first finding underscores the crucial role of intra-individual communication, revealing that an advantage in this form of communication is one of essential factors to counteract the invasion of cheaters who refrain from emitting BVOCs.

The second finding is also related to the persistence of the BVOC emission strategy against cheaters. The limited spatial scale of BVOC signaling was found to be effective against invasion by cheaters because access to public goods is limited to cheaters. Short range of seed dispersal was also effective in maintaining BVOC emission because clustering of individuals with the BVOC emission strategy allows cooperators to interact with each other preferentially and favor cooperation (Chao and Levin, 1981; Fletcher and Doebeli, 2008; Nowak and May, 1992). The valid spatial scale of BVOC signaling could depend on BVOC profiles, reactivity, and plant species (Masui et al., 2023). Previous studies reported a limited range of BVOC signaling, ranging from 0.6m in sagebrush to approximately 5 m in the Japanese beech (Hagiwara et al., 2021; Karban et al., 2006), which could contribute to the maintaining BVOC emission. More research is needed to explore and estimate the spatial scale of BVOC signaling in diverse plant species.

The third finding highlights the possibility that variations in BVOC emission levels, such as high and low BVOC emission within a population, can be maintained as an evolutionarily stable state. Empirical studies in various plant species have reported that the amount of BVOC emission varies among plant populations. The compositional ratio and emission rates of terpenoids varied between genetically differentiated local populations of Japanese cedar, *Cryptomeria japonica* (Hiura et al., 2021), and the total amounts of released BVOCs altered among the accessions of lima bean, *Phaseolus lunatus* (Ballhorn et al., 2008). In sagebrush, chemotypes were highly heritable (Karban et al., 2014), and each population comprised different compositions of the chemotypes determined by the dominant compounds of BVOC emission profiles (Grof-Tisza et al., 2021). The value of releasing BVOCs for plants to induce resistance can change (Heil and Baldwin, 2002) if biotic interactions harming plants (e.g., herbivory) can decrease the fitness of plants (Stowe et al., 2013; Syller and Grupa, 2016). While studies have focused on the variation in BVOC emission levels among plant populations with relatively different conditions for the environment, genetic background, and biotic interactions, fewer studies have focused on the variation within populations. Although ecological implications for the variation in BVOC emission levels within populations are unclear, our results show that the BVOC emission strategy is relatively tolerant of cheaters using BVOCs as public goods without production costs. In short, such variation could be explained by tolerant coexistence with cheaters. In addition, there are also qualitative variations in BVOC emission as shown in the above studies. We assumed that the perception of a greater amount of BVOCs would induce strong resistance to simplify models and omitted the effect of BVOC type on signaling. The use of various BVOC signals is intricately regulated by the biosynthetic pathways depending on the availability of carbon, nitrogen, sulfur, and energy (Dudareva et al., 2013; Rasheed et al., 2023), so the available type and amount of BVOCs are limited and they can affect the efficiency and accuracy of information transfer of plants. Since various plants release a mixture of BVOCs and transfer information, future theoretical studies may need to focus on qualitative differences in BVOC emissions as well.

The future accumulation of knowledges on BVOC types, emissions, and their seasonality across a wider range of species and populations will serve to validate the predictions made in this theoretical study and offer a comprehensive understanding of the evolution of BVOC emission strategies.

## Author Contributions

### Sotaro Hirose

Data curation (lead); investigation (lead); methodology (equal); software (lead); visualization (lead); writing – original draft (lead); writing – review and editing (equal). **Akiko Satake**: Funding acquisition (lead); investigation (supporting); methodology (equal); supervision (lead); visualization (supporting); writing – review and editing (equal).

## Acknowledgements

The authors would like to thank to Tomika Hagiwara for her valuable comments on our initial draft. This work was done in support of JSPS KAKENHI Grant Number JP23H04966 and JP23H04965 to A. Satake.

## Competing Interests Statement

The authors have no competing interests.

## Data Availability Statement

Relevant data and code are available at Zenodo https://doi.org/10.5281/zenodo.11112818 (Hirose and Satake, 2024).

## Appendix

### Appendix A. Parameter dependence in the random distribution model: herbivory rate and cost of BVOC production

We developed the random distribution model and analyzed the influence of herbivory rate *p* and cost of BVOC production *c* (Fig. A1). Parameter areas where BVOC emitters dominate or coexist with non-emitters decrease when the value of *c* is large or that of *p* is small (Fig. A1). Therefore, the emission strategy is less likely to evolve when the cost of BVOC production is high due to the small benefits of the emission and when the herbivory rate is low due to the small benefits of releasing BVOCs to suppress the damage. Based on these intuitive results, we analyzed the influence of the intra- and inter-individual communication effects under conditions that the BVOC emission strategy is more likely to evolve: the BVOC production cost is low and the herbivory rate is high (Fig. 2-4, A2-A5).

### Appendix B. Influence of the initial value in lattice model

We developed the lattice model and ran simulations by giving three patterns of initial values for each parameter set that consists of the intra- and inter-communication effect. As with the random distribution model, we could not find apparent changes in the frequency depending on the initial values such as the bistability (Fig. A2-A5).

**Figure A1:**
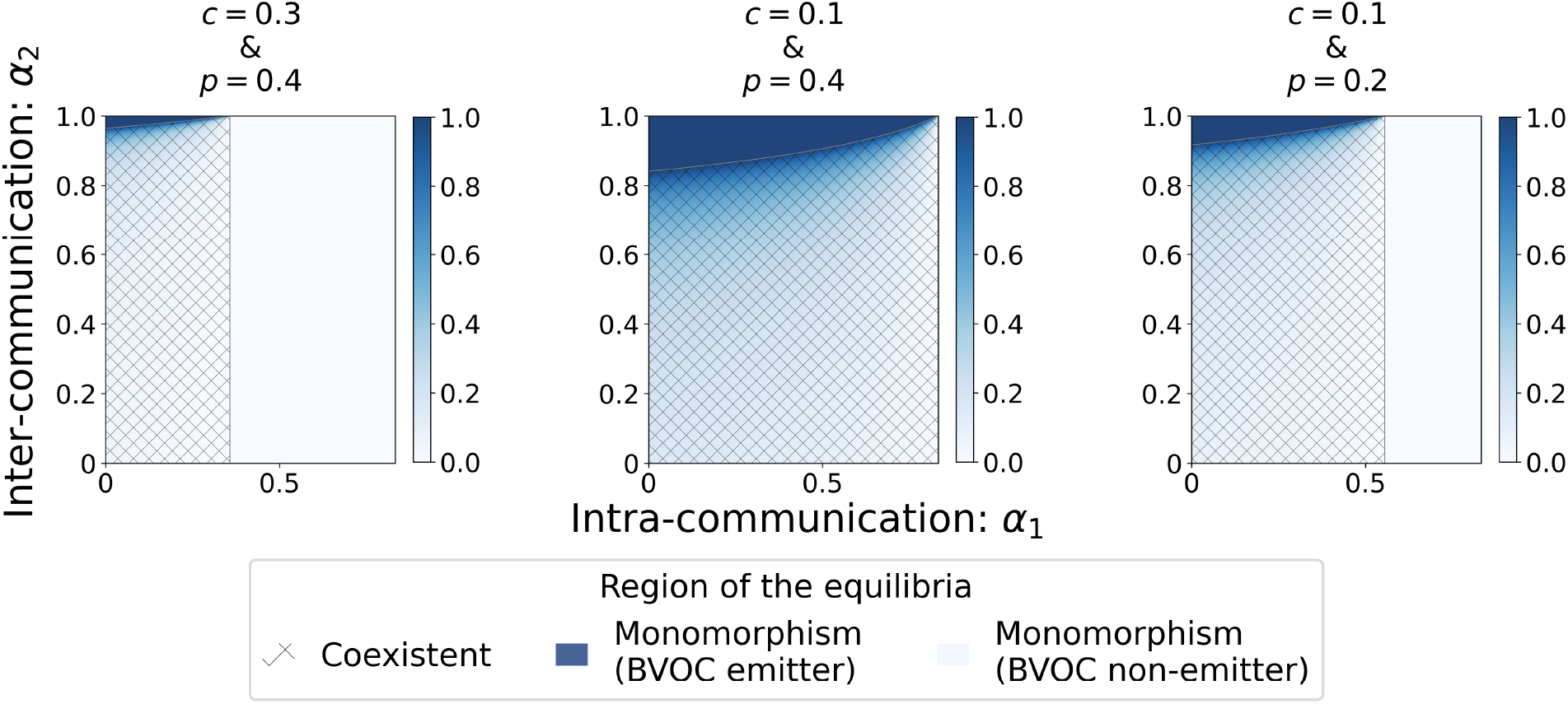
Effects of herbivory rate and cost of BVOC production on the evolutionary consequences of the BVOC emission strategy. Each panel shows the values of the local stable equilibria (the frequency of emitters) on the *α*_1_-*α*_2_ plane in the random distribution model. The vertical axis corresponds to the inter-individual communication effect and the horizontal axis to the intra-individual communication effect. The parameter area can be divided into three types: The shaded area has a stable equilibrium where BVOC emitters coexist with non-emitters, the dark blue one has a stable equilibrium where emitters are dominant, and the white one has a stable equilibrium where non-emitters are dominant. Parameters are as follows: *c* = 0.3, *p* = 0.4, *n* = 8 (left panel); *c* = 0.1, *p* = 0.4, *n* = 8 (central panel); *c* = 0.1, *p* = 0.2, *n* = 8 (right panel).

**Figure A2:**
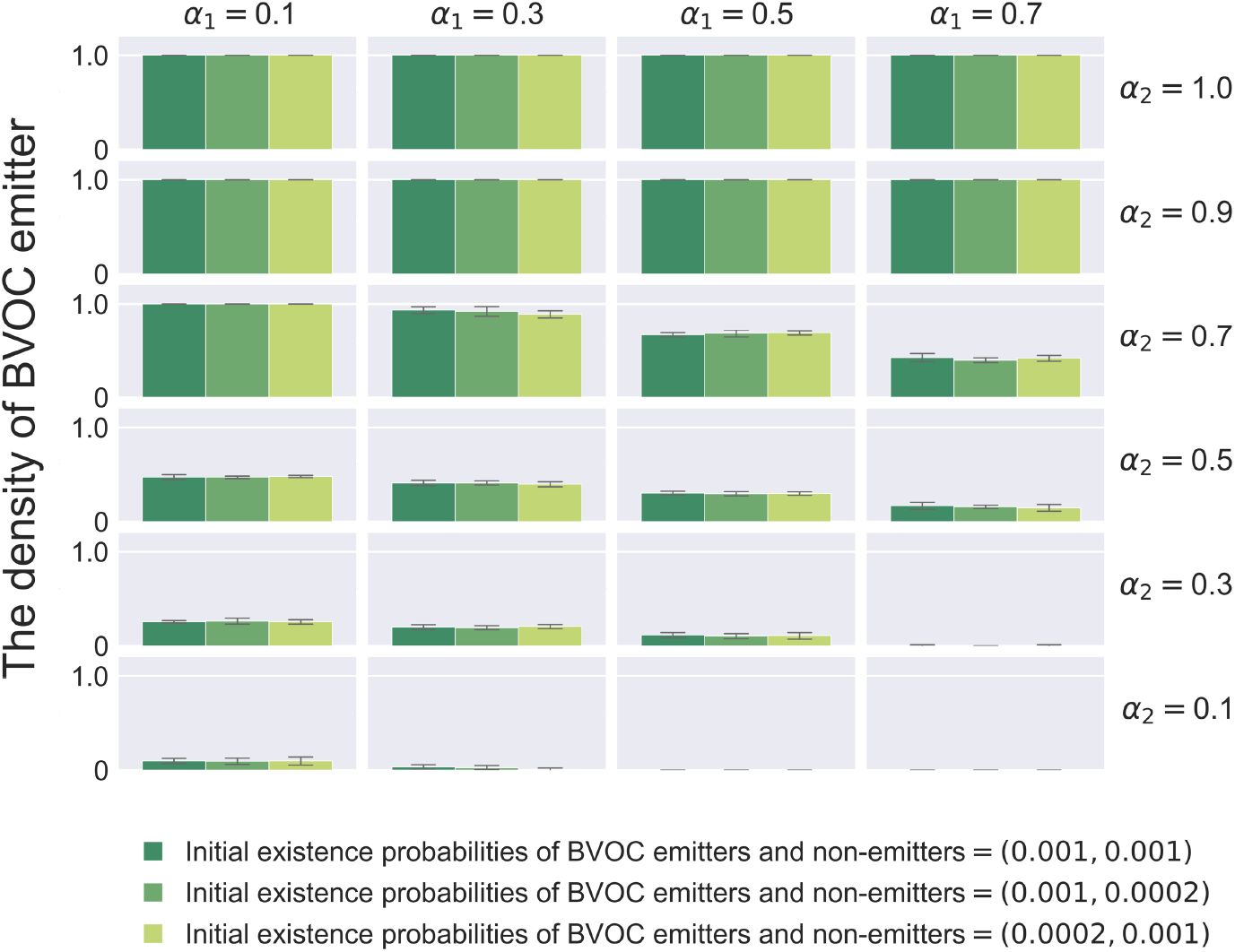
Effects of the initial number of emitters and non-emitters on the evolutionary consequences of the BVOC emission strategy. Each panel shows three patterns of the average density of BVOC emitter obtained from ten times simulations at thirty thousand steps in the lattice model. Initial existence probabilities of BVOC emitters and non-emitters in the lattice structure is as follows: (0.001, 0.001; dark green); (0.001, 0.0002; green); (0.0002, 0.001; light green). Parameters are *c* = 0.1, *p* = 0.4, *d* = 0.0*D, r*^*BVOC*^ = 1, *r*^*D*^ = 1.

**Figure A3:**
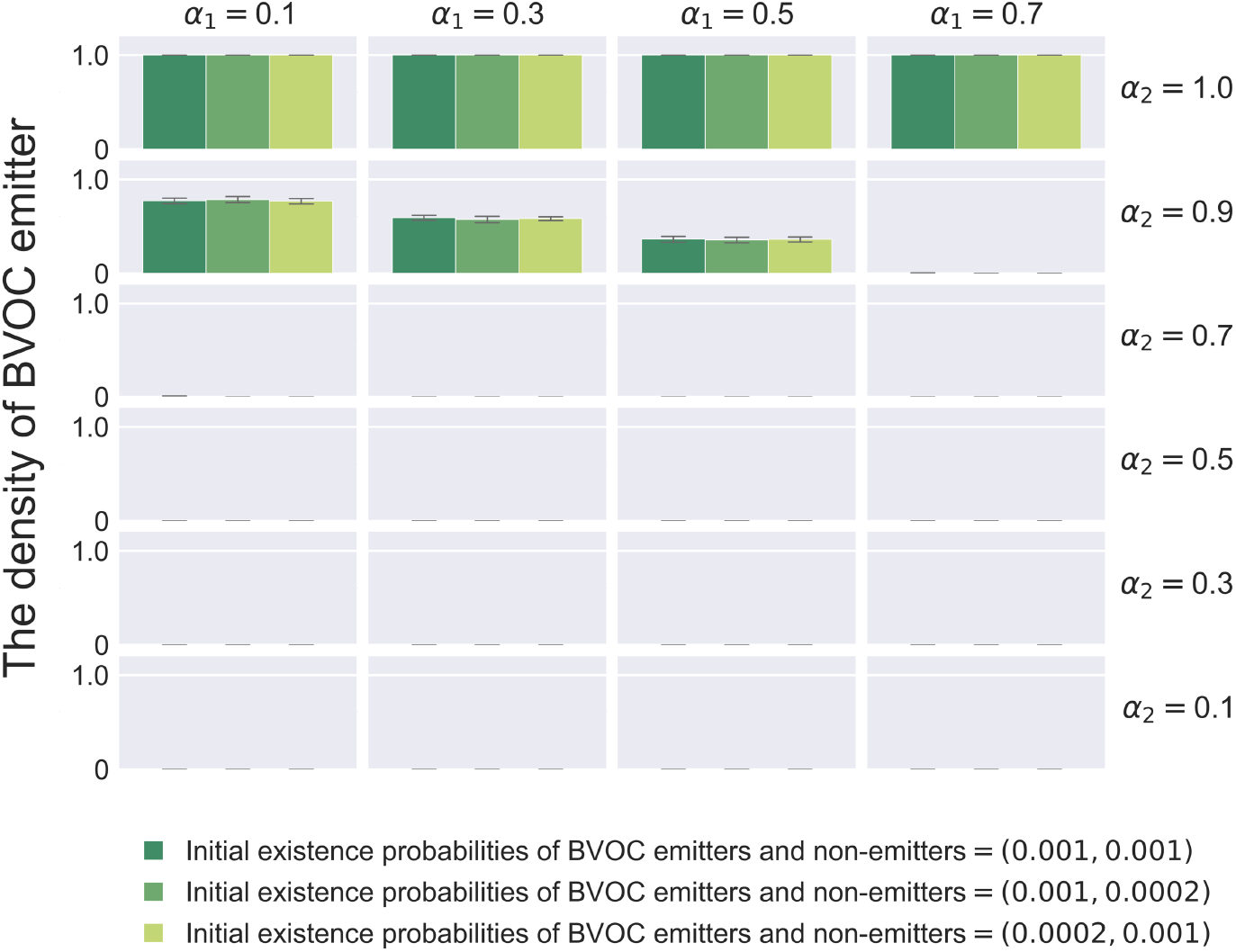
Effects of the initial number of emitters and non-emitters on the evolutionary consequences of the BVOC emission strategy. Each panel shows three patterns of the average density of BVOC emitter obtained from ten times simulations at thirty thousand steps in the lattice model. Initial existence probabilities of BVOC emitters and non-emitters in the lattice structure is as follows: (0.001, 0.001; dark green); (0.001, 0.0002; green); (0.0002, 0.001; light green). Parameters are *c* = 0.1, *p* = 0.4, *d* = 0.0*D, r*^*BVOC*^ = 2, *r*^*D*^ = 1.

**Figure A4:**
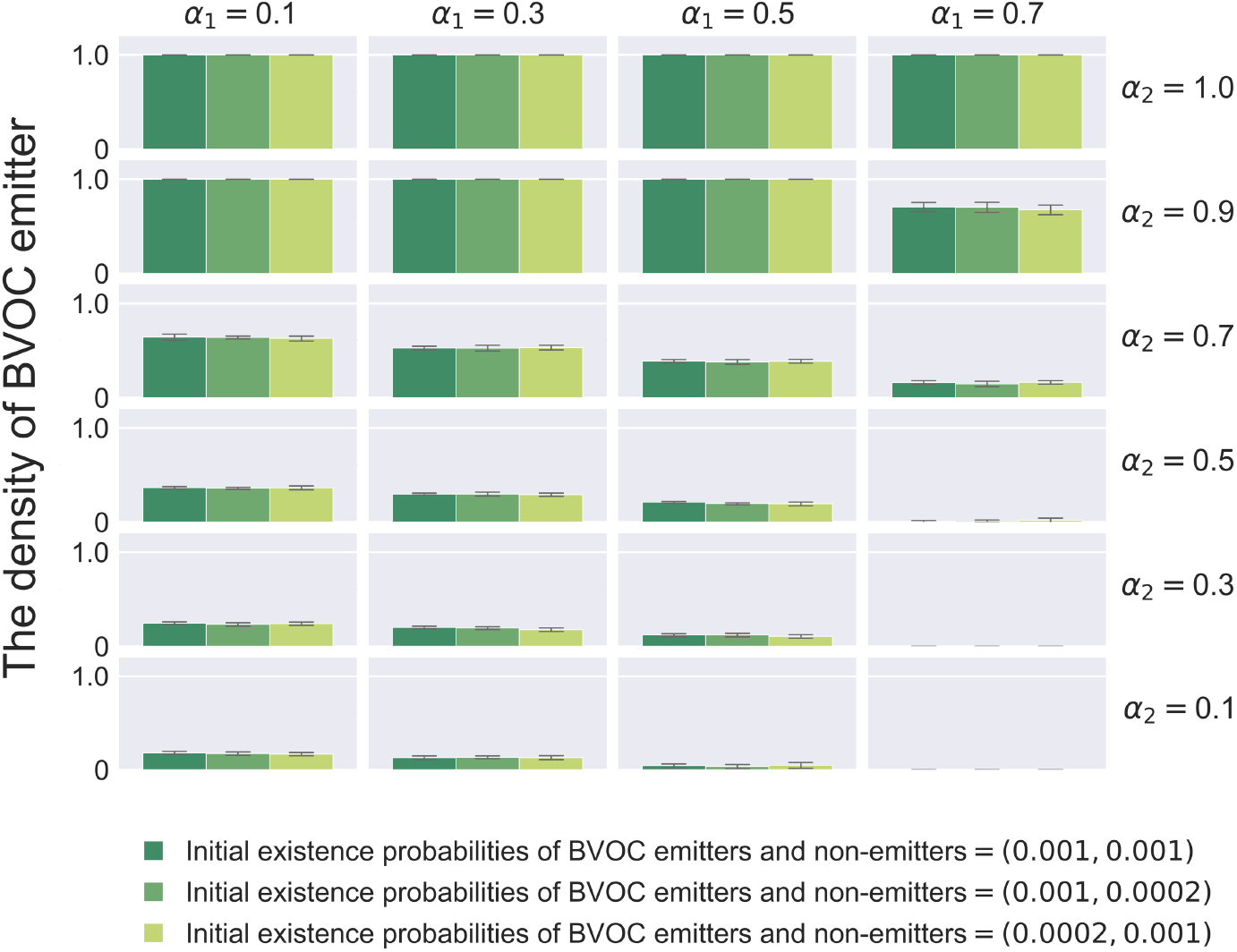
Effects of the initial number of emitters and non-emitters on the evolutionary consequences of the BVOC emission strategy. Each panel shows three patterns of the average density of BVOC emitter obtained from ten times simulations at thirty thousand steps in the lattice model. Initial existence probabilities of BVOC emitters and non-emitters in the lattice structure is as follows: (0.001, 0.001; dark green); (0.001, 0.0002; green); (0.0002, 0.001; light green). Parameters are *c* = 0.1, *p* = 0.4, *d* = 0.0*D, r*^*BVOC*^ = 1, *r*^*D*^ = 2.

**Figure A5:**
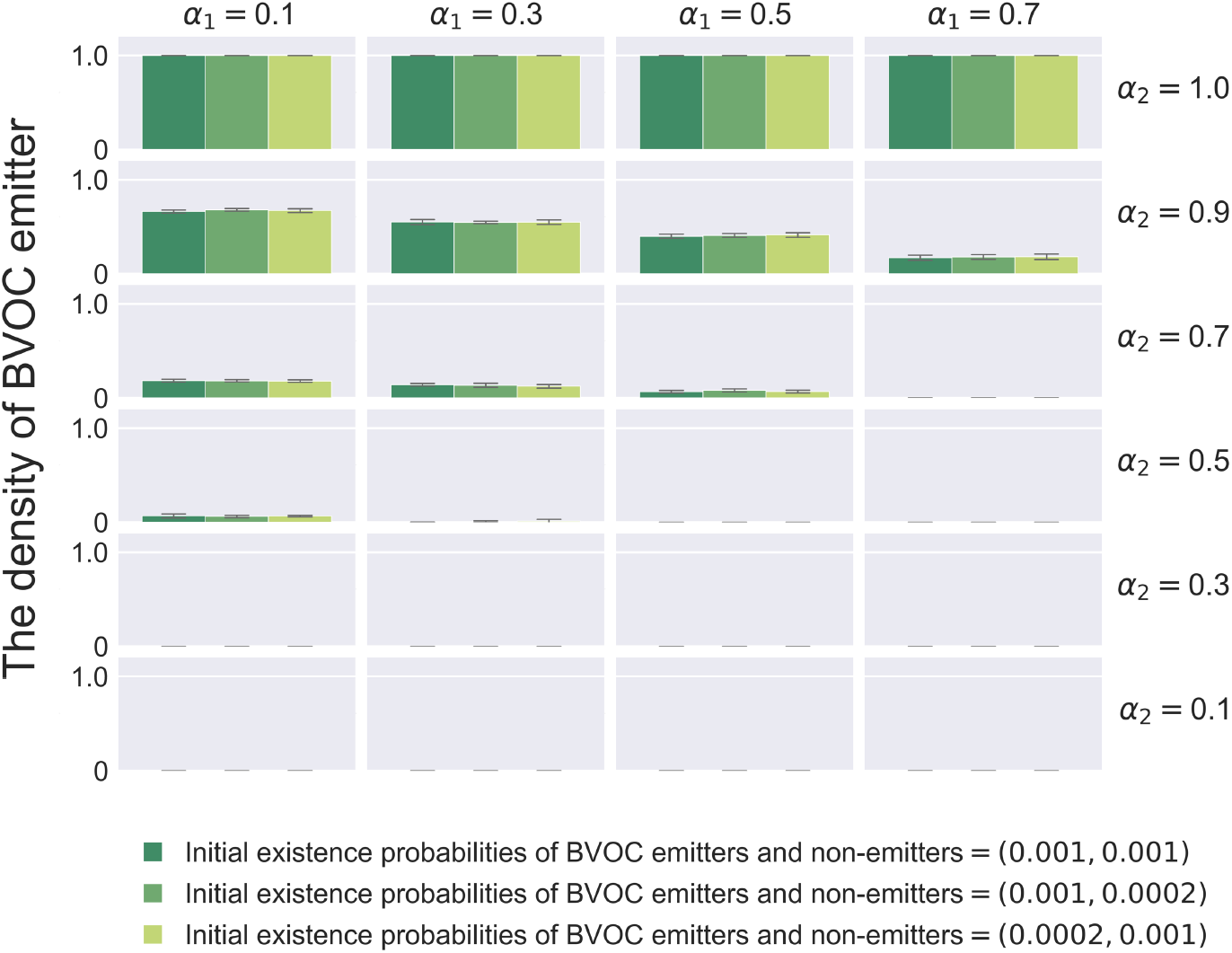
Effects of the initial number of emitters and non-emitters on the evolutionary consequences of the BVOC emission strategy. Each panel shows three patterns of the average density of BVOC emitter obtained from ten times simulations at thirty thousand steps in the lattice model. Initial existence probabilities of BVOC emitters and non-emitters in the lattice structure is as follows: (0.001, 0.001; dark green); (0.001, 0.0002; green); (0.0002, 0.001; light green). Parameters are *c* = 0.1, *p* = 0.4, *d* = 0.0*D, r*^*BVOC*^ = 2, *r*^*D*^ = 2.

## Notes

https://doi.org/10.5281/zenodo.11112818

### Competing Interest Statement

The authors have declared no competing interest.

